# Global genomic diversity of *Pseudomonas aeruginosa* in bronchiectasis

**DOI:** 10.1101/2024.01.30.577916

**Authors:** N.E. Harrington, A. Kottara, K. Cagney, M.J. Shepherd, E.M. Grimsey, T. Fu, R.C. Hull, D.Z. Childs, J.L. Fothergill, J.D. Chalmers, M.A. Brockhurst, S. Paterson

## Abstract

**Background:** *Pseudomonas aeruginosa* is the dominant pathogen causing lung infections in people with both cystic fibrosis (CF) and bronchiectasis, associated with poorer outcomes. Unlike CF, bronchiectasis has been a neglected disease. More extensive genomic studies of larger bronchiectasis patient cohorts and within patient sampling are needed to improve understanding of the evolutionary mechanisms underpinning *P. aeruginosa* infections to guide novel and improved treatments.

**Methods:** We have performed genome sequencing of 2,854 *P. aeruginosa* isolates from 180 patients attending clinics worldwide to analyse the genomic diversity between and within patient infections.

**Results:** We observed high genetic diversity between infections with low incidence of highly transmissible strains. Our genomic data provide evidence for the mutational targets driving *P. aeruginosa* evolution in bronchiectasis. Some functions found to gain mutations were comparable to CF, including biofilm and iron acquisition, whilst others highlighted distinct evolutionary paths in bronchiectasis such as pyocin production and resistance, and a novel efflux pump gene (PA1874). We also show a high incidence of antimicrobial resistance-associated mutations and acquired resistance genes, in particular multidrug efflux and fluoroquinolone resistance mechanisms.

**Conclusions:** Our findings highlight important differences between *P. aeruginosa* infections in bronchiectasis and CF and provide evidence of the relatively minor role transmissible strains play in bronchiectasis. Our study provides a 10-fold increase in the available genomic data for these infections and is a global resource to improve our knowledge and understanding, to facilitate better patient outcomes.

**Summary:** The largest genomic study of *Pseudomonas aeruginosa* bronchiectasis isolates to-date, providing an unprecedented global genomic resource. We highlight important differences between bronchiectasis and cystic fibrosis, including key genes under selection.

## Introduction

Bronchiectasis is a chronic respiratory disease where people suffer ongoing symptoms of cough, sputum production and frequent respiratory infections, leading to progressive lung function decline and reduced quality of life^1–3^. The disease is defined by abnormal, permanent dilation of the bronchi resulting in impaired clearance of mucus from the airways. The impaired mucocillary clearance leads to chronic infection and inflammation, which further exacerbates lung damage, establishing a progressive, vicious cycle^4^. In contrast to cystic fibrosis (CF), a rare genetic cause of bronchiectasis caused by mutations in the cystic fibrosis transmembrane conductance regulator (CFTR) gene, most cases of non-CF bronchiectasis (hereafter referred to as bronchiectasis) are of unknown cause and are not monogenic^5^.

Over time, the lungs of most people with bronchiectasis become chronically infected with bacterial pathogens, predominantly *Pseudomonas aeruginosa*. In a recent analysis of the European bronchiectasis registry, involving 16,963 patients from 28 countries, *P. aeruginosa* infection was identified in ~25% of all patients^6^. However, there was marked regional variation, with more than 50% of patients infected with *P. aeruginosa* being in Southern European countries, and lower rates observed in Northern Europe. Outside of Europe, *P. aeruginosa* has been shown to be the dominant pathogen in the United States^7^, China^8^, India and Australia^9^. The presence of *P. aeruginosa* infection in people with bronchiectasis is associated with poorer outcomes, and patients with *P. aeruginosa* infection have been found to have a nearly 7-fold higher risk of hospitalisation and 3-fold increased risk of mortality compared to patients without^10^.

Bronchiectasis has been a neglected disease with limited research, consequently there are far fewer genomic studies characterising the associated *P. aeruginosa* infections than in CF. To date there have been only two single-country genomic epidemiology studies solely focused on *P. aeruginosa* in bronchiectasis. A recent investigation of 130 genomes isolated from 110 adult patients attending a single bronchiectasis clinic in Germany^11^ suggested that bronchiectasis infections are caused by diverse sequence types (STs), representative of the global diversity of *P. aeruginosa*. The incidence of multiple patients with an infection caused by the same ST was rare. Similarly, a study of 189 *P. aeruginosa* genomes from 91 adult patients attending 16 clinics in the UK^12^ showed a high diversity of STs. This study also reported ~30% of patients for whom multiple *P. aeruginosa* genomes were obtained that had mixed ST infections, suggesting multiple acquisition events. As mixed ST *P. aeruginosa* infections have been shown to more readily acquire antimicrobial resistance (AMR)^13^, this could pose a clinical risk to patient health and urgently requires further study.

In CF, *P. aeruginosa* undergoes a characteristic suite of genomic changes enabling adaptation to the lung environment. This process drives extensive genetic diversification of the infecting population through mutations commonly affecting traits such as AMR, biofilm formation and motility, and regulatory systems controlling a range of functions including virulence factor production. These genomic changes are believed to directly impact patient health, by prolonging infection and reducing the effectiveness of antimicrobial treatments^14,15^.

Whether such evolutionary diversification occurs in bronchiectasis *P. aeruginosa* infections is unclear due to the extremely limited data^12^. Given the more variable aetiology, disease phenotypes, and treatment responses of bronchiectasis compared to CF^16^, it is probable that the evolutionary pathways of bronchiectasis infections are less predictable than for CF infections. More extensive genomic studies that sample the within patient diversity of *P. aeruginosa*, across a larger number of bronchiectasis patients, have been urgently required to better understand the evolutionary mechanisms operating in bronchiectasis infections to guide improved treatment.

We have used genome sequencing to analyse a unique global collection of 2,854 *P. aeruginosa* bronchiectasis isolates, obtained during the ORBIT3 Phase III clinical trial for inhaled liposomal ciprofloxacin^17^. Here we analyse the genomic diversity of *P. aeruginosa* within and between 180 patient infections sampled at baseline prior to antibiotic treatment.

## Materials and Methods

### *Pseudomonas aeruginosa* isolation and growth

Sputum samples were collected from a cohort of 180 patients with bronchiectasis at the start of the ORBIT3 clinical trial^17^, and stored at -80 °C. To isolate *P. aeruginosa* from the sputum, an equal volume of Sputasol solution (SR0233, Oxoid) was added to each sample, mixed until liquefaction was complete, and then incubated in a 37 °C shaking incubator (240 rpm) for 30 min. Subsequently, 100 μl was spread onto Cetrimide Agar (22470, NutriSelect® Plus) plates and incubated in a static incubator at 37 °C for 24 -48 h. These populations were further streaked onto Cetrimide Agar plates and incubated in a static incubator at 37°C for 24 – 48 h, and *P. aeruginosa* colonies were then isolated from each population. These selected colonies were grown in King’s B medium consisting of 20 g l^-1^ Bacto proteose peptone No.3 (Gibco), 1.5 g l^-1^ Potassium phosphate dibasic trihydrate (P5504, Sigma), 1.5 g l^-1^ Magnesium sulfate heptahydrate (M1880, Sigma) and 10 g l^-1^ Glycerol (49770, Honeywell) in a static incubator at 37 °C for 48 h. The presence of *P. aeruginosa* was confirmed using polymerase chain reaction (PCR) and primers targeting the species-specific 16S rRNA gene of *P. aeruginosa*^18^. The primer sequences used were: forward, 5’ - GGG GGA TCT TCG GAC CTC A - 3’; reverse, 5’ - TCC TTA GAG TGC CCA CCC G - 3’. The PCR protocol included the following thermocycling program: initial denaturation at 95 °C for 5 min, followed by 35 cycles of denaturation at 95 °C for 20 s, annealing at 58 °C for 20 s, extension at 72 °C for 40 s, and a final extension at 72 °C for 10 min.

### DNA extraction and sequencing

Isolates for sequencing were cherrypicked using an Opentrons OT-2 Liquid Handler and DNA extraction was then performed using the Quick-DNA Fecal/Soil Microbe Kit (Zymo Research) on a KingFisher Flex instrument (Thermo Fisher Scientific), following the manufacturers protocol. Eluted DNA was quantified using a Qubit Flex Fluorometer (Thermo Fisher Scientific) and samples kept at -80 °C for storage. Samples were then normalized to 7.7 ng μl^-1^ using a MANTIS Liquid Handler (Formulatrix) and library preparation performed for 20 ng of input DNA using NEBNext Ultra II FS DNA Library Prep Kit for Illumina (New England Biosciences) on a Mosquito HV liquid handling instrument (SPT Labtech), following the manufacturers protocol, miniaturized to 0.1 recommended volume. Library fragments were then amplified and 10 bp index sequences (Integrated DNA Technologies) were incorporated using polymerase chain reaction (PCR).

Following library preparation, the libraries were quantified using a Qubit Flex Fluorometer and size analysis was performed on a Fragment Analyser (Agilent). The average library length and concentration of each library was used to pool the libraries in an equimolar manner using a Mosquito X1 (SPTLabtech). The final pooled libraries cleaned and concentrated using AMPure XP beads (Beckman Coulter) to remove any remaining adaptor sequences. The average length of the final pool of libraries was analysed using a Bioanalyzer (Agilent) and the concentration was quantified using a Qubit fluorometer. Samples were then sequenced on an Illumina NovaSeq 6000 150 bp paired-end run by the Centre for Genomics Research, University of Liverpool, who then trimmed the raw fastq files using Cutadapt v1.2.1^19^.

### Assembly, annotation and MLST analysis

All reads were then quality checked with FastQC v0.11.9^20^, and *de novo* assembly performed using Unicycler v0.5.0^21^. Each genome was annotated with Bakta v1.6.0^22^, and quality checked using the quality control (QC) pre-processing script as part of Panaroo v1.2.10^23^, QUAST v5.2.0^24^ and Busco v5.4.3^25^, and any isolates with poor assemblies were removed. Panaroo QC was also used to detect samples contaminated with reads from other species (indicative of initial sample contamination) or non-*P. aeruginosa* isolates, which were then removed from the analyses. All remaining assemblies were sequence typed with mlst v2.11^26^, using the PubMLST database (https://pubmlst.org/)^27^.

### Pangenome and phylogeny

The pangenome of each sample group of interest: all isolates, each phylogroup and individual patients, was constructed using Panaroo v1.2.10^28^ with mafft alignment. The core genome produced was then used to determine the core SNP phylogeny for each group using SNP-sites v2.5.1^29^. Maximum likelihood phylogenetic trees were then constructed based on these using IQ-Tree v2.0.0.3^30^ (1000 bootstraps) with ModelFinder for model selection^31^ and ascertainment bias correction.

### Prophage and plasmid detection

To screen for viral DNA and detect prophage regions in each genome, VirSorter v1.0.5^32^ was used. Viral sequences detected in each assembled genome assigned to prophage category 1 (#4) and prophage category 2 (#5) were considered prophage regions, selected at random for confirmation using PHASTEST^33–35^. To detect possible plasmids in assemblies, Abricate v0.7^36^ was used to identify any plasmid replicons against the PlasmidFinder database^37^.

### Mutation diversity

To investigate mutational diversity, all SNPs in each genome were first identified against the annotation file for the closest reference strain for each phylogenetic group (group 1: *P. aeruginosa* PAO1, group 2: PA14, group 3+: PA7) using Snippy v4.6.0^38^. The ‘snippy-core’ script, part of Snippy, was then used to determine core SNPs amongst the different groups of isolates described. This identified all SNPs in core sites amongst each group of isolates (e.g. phylogenetic groups/patient populations). SNP pairwise distance matrices were then produced using snp-dists v0.8.2 and visualised as heatmaps using the *pheatmap* package in R v4.3.1^39^. SnpEff v5.0e^40^ was subsequently used to annotate the core SNPs identified. Comparisons to determine polymorphic genes and fixed mutations were performed using these outputs in R. Ancestral state reconstruction, under an equal-rates model, of loss of function core SNPs observed in multiple isolates was performed and mapped to the phylogeny using SIMMAP^41^ and the *phytools* package^42^ in R v4.3.1.

### Antimicrobial resistance

To detect the presence of antimicrobial resistance genes in each genome assembly, ResFinder 4.0^43^ was used. There was one isolate from a patient where OXA-50, expected in all *P. aeruginosa* genomes, was not detected by ResFinder, however visual inspection using IGV v2.16.1^44^ confirmed this was a false negative. To detect AMR-associated mutations, RGI was used against the Comprehensive Antibiotic Resistance Database (CARD)^45^.

### Reference strains and gene information

Four reference strain genomes were included in this work: *P. aeruginosa* PAO1 (NCBI, GCF_000006765.1), *P. aeruginosa* PA14 (NCBI, GCF_000014625.1), *P. aeruginosa* LESB58 (NCBI, GCF_000026645.1) and *P. aeruginosa* PA7 (NCBI, GCF_000017205.1). All gene information was sourced from pseudomonas.com^46^.

### Statistical analysis

All statistical analyses were performed using R v4.3.1. The *pegas* package^47^ was used to conduct AMOVAs (1000 permutations), and to calculate nucleotide diversity (π) alongside the *vcfR* package^48^.

### Data availability

All sequencing data (reads and assemblies) are available at the European Nucleotide Archive (ENA) (accession number: PRJEB65845).

## Results

We sequenced 2,854 *P. aeruginosa* isolates from 180 patients, representative of the bronchiectasis demographic (~16 isolates per patient; Fig S1), and performed *de novo* assembly (median number of contigs = 80; see Table S1) and annotation for each genome. A phylogeny of all isolates and the common reference strains PAO1, PA14, LESB58 and PA7 was constructed based on single nucleotide polymorphisms (SNPs) in the 4,344 core genes; 83% of isolates belonged to phylogenetic group 1 (PAO1-like), and 14% belonged to phylogenetic group 2 (PA14-like) (Table S2). The remaining 3% of isolates from 5 patients branched distantly from group 1 and group 2 isolates, closest to the reference strain PA7 (Fig 1A) and were classified within group 4 or 5^49^. We observed strong phylogenetic clustering of isolates by patient, but there was no clear evidence of clustering driven by geographic region (Fig 1B). Most patients (77%) contained a single, distinct ST (Fig S2). Only 18 patients had infections caused by clonal STs, most commonly Clone C, and CF epidemic STs were rare (6 patients; Table S3). There were only 3 patients (2%) that had mixed ST infections (Fig S2). Together these data suggest that transmissible strains and superinfection by multiple strains do not play a major role in bronchiectasis.

**Figure 1.**
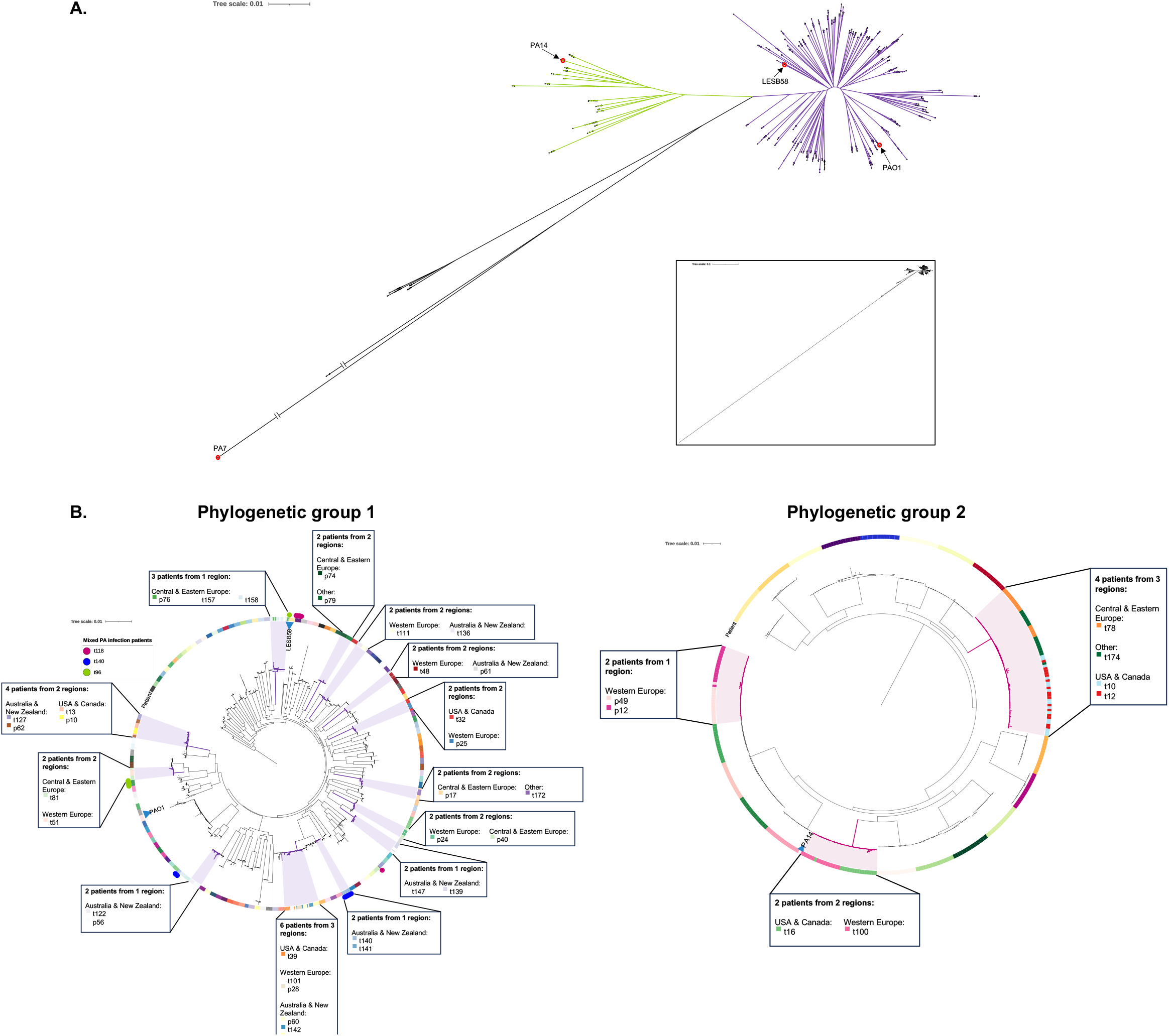
Core single nucleotide polymorphism (SNP)-based phylogenetic trees of *Pseudomonas aeruginosa* isolates from people with bronchiectasis and 4 reference strains: PAO1, LESB58, PA14 and PA7. **(A)** Phylogenetic tree including all isolates and reference strains, each group is denoted by coloured branches (group 1: right, purple, group 2: left, green, group 3+: down, black), and the reference strains are circled and labelled. **(B)** Phylogenetic trees of the two dominant phylogenetic groups, group 1 (PAO1-like) and group 2 (PA14-like). The coloured rings show the patient each isolate was taken from, and the circles in the outer ring on the group 1 tree show the possible mixed infection isolates (see legend). The labelled boxes and highlighted regions show clusters of patients whose isolates branched together and detail the clinic region these patients attended.

Group 2 isolates had higher nucleotide diversity than group 1 isolates (group 2 π=6.11x10^-3^; average 1 SNP per 164 bases; group 1 π=2.93x10^-3^; average 1 SNP per 342 bases) and genetic diversity between patients far exceeded that observed within patients, accounting for 99.9% of total genetic diversity in both group 1 (df = 145, σ < 0.01, *P* < 0.01) and group 2 (df = 25, σ < 0.01, *P* < 0.01). Similarly, gene content varied extensively in both phylogenetic groups (group 1: 4,580 core genes and 14,138 accessory genes, n=2,349 isolates; group 2: 4,627 core genes and 6,817 accessory genes, n=415 isolates) and showed greater variation between than within patients (1-sample t-tests: group 1 *t*_145_= -725.53, *P* < 0.01; group 2 *t*_25_= - 202.15, *P* < 0.01; Fig 2). Nonetheless, we did observe strong evidence supporting within-patient diversification. Focusing on single ST infections, where genetic diversity can be assumed to have arisen by mutation *in situ*, average pairwise core SNP distance between co-existing isolates per patient ranged from 3.86x10^-5^ (1 SNP per 25,926 bases) to 2.81x10^-7^ (1 SNP per 3,555,101 bases) for group 1, and from 2.08x10^-5^ (1 SNP per 48,045 bases) to 1.95x10^-7^ (1 SNP per 5,127,358 bases) for group 2 (Fig S3), and did not differ between the phylogenetic groups (T test: *t*_45.04_ = 0.26, *P* = 0.80). In some patients, elevated within-patient genetic diversity was associated with mutations in genes encoding DNA mismatch repair (MMR) or break excision repair (BER) likely to cause hypermutation (Fig S3). Mobile genetic elements also played a role generating within patient diversity: Prophage regions (20 to 433 genes in size; mean = 135 genes) were observed in all isolates (range 1 to 9 regions per genome; mean = 3; Table S4), and coexisting isolates from the same patient often varied in their prophage number (50%; n = 89 patients) and/or the gene content of prophage regions (86%; n = 153 patients). Plasmids were comparatively much rarer across all isolates, detected in only 53 isolates from 6 patients, but variable plasmid carriage between coexisting isolates caused the highest level of within patient gene content variation observed.

**Figure 2.**
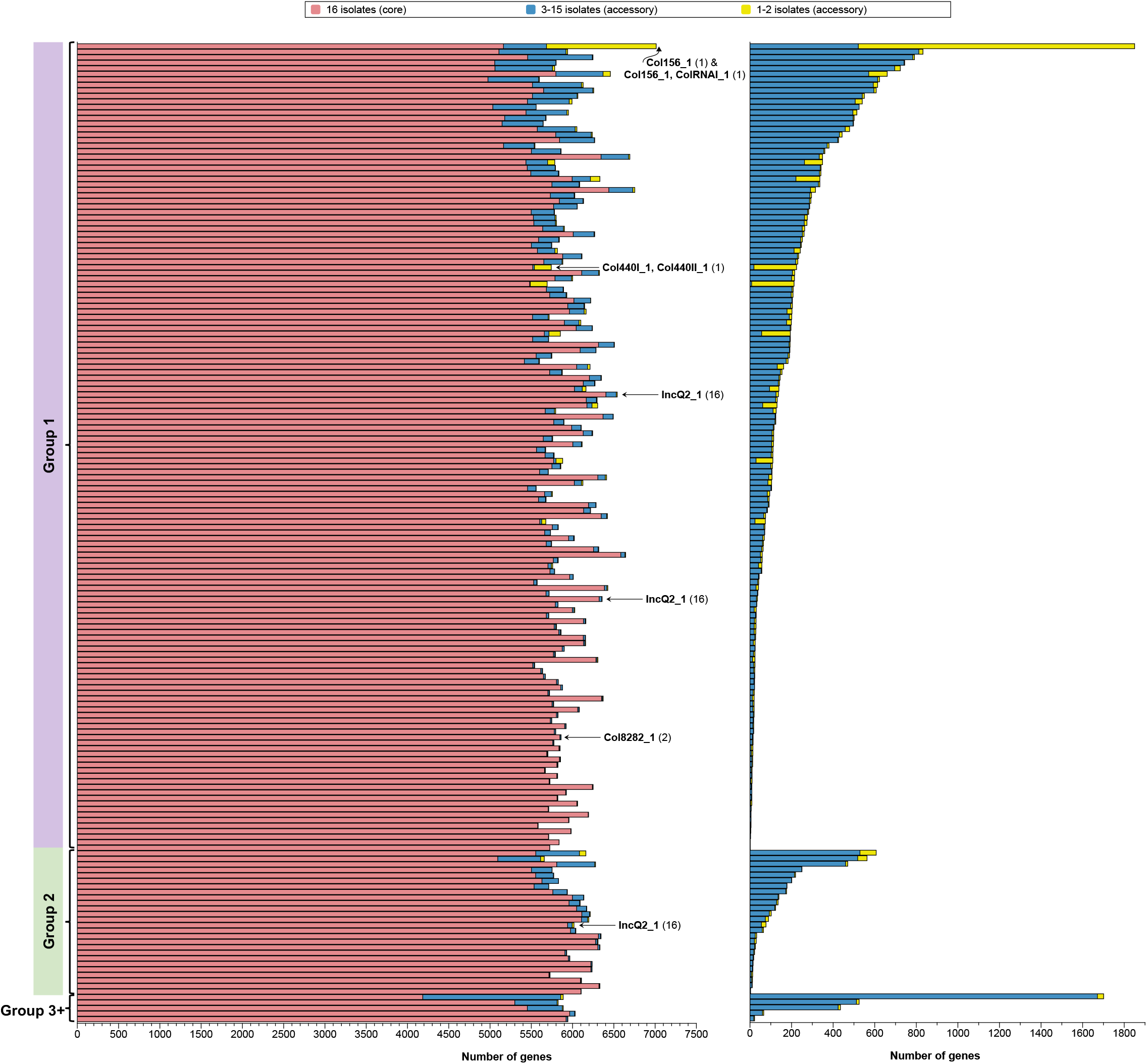
**(A)** The number of genes in the core and accessory *Pseudomonas aeruginosa* genome in sequenced populations from people with bronchiectasis. **(B)** The number of genes in the accessory genome of each population, in the same patient order as graph A. In both graphs, each bar represents a patient and ordered by phylogenetic group. The patients with plasmids detected are labelled with the replicon and the number in brackets indicates how many sequenced isolates carried the replicon/s from that patient.

Putative targets of diversifying selection within patients were determined by identifying genes that harboured non-synonymous polymorphisms across multiple patients. Polymorphic genes (Table S5) identified in both group 1 and group 2 (Fig 3) included a novel efflux pump gene (PA1874), sigma factor-encoding gene *algU* (PA0762) and the flagellar-associated gene *flgK* (PA1086). Additionally, in group 1, polymorphic genes associated with a range of key functions were identified, including motility (*fliC, chpA* and *pilB*), cell envelope (*oprE, opmH, oprD, pagL, migA*), alginate production (*mucA, algG*) and iron acquisition (*fptA, pvdL, pvdS, pchE, hemA, pvdP*), as well as *pdtA*, encoding a filamentous haemagluttinin linked with adhesion and virulence. For group 2, the most common polymorphic gene was PA1572 (Fig 3B), also known as *nadK1*, which encodes an ATP-NAD kinase associated with response to reactive oxygen species^50^.

**Figure 3.**
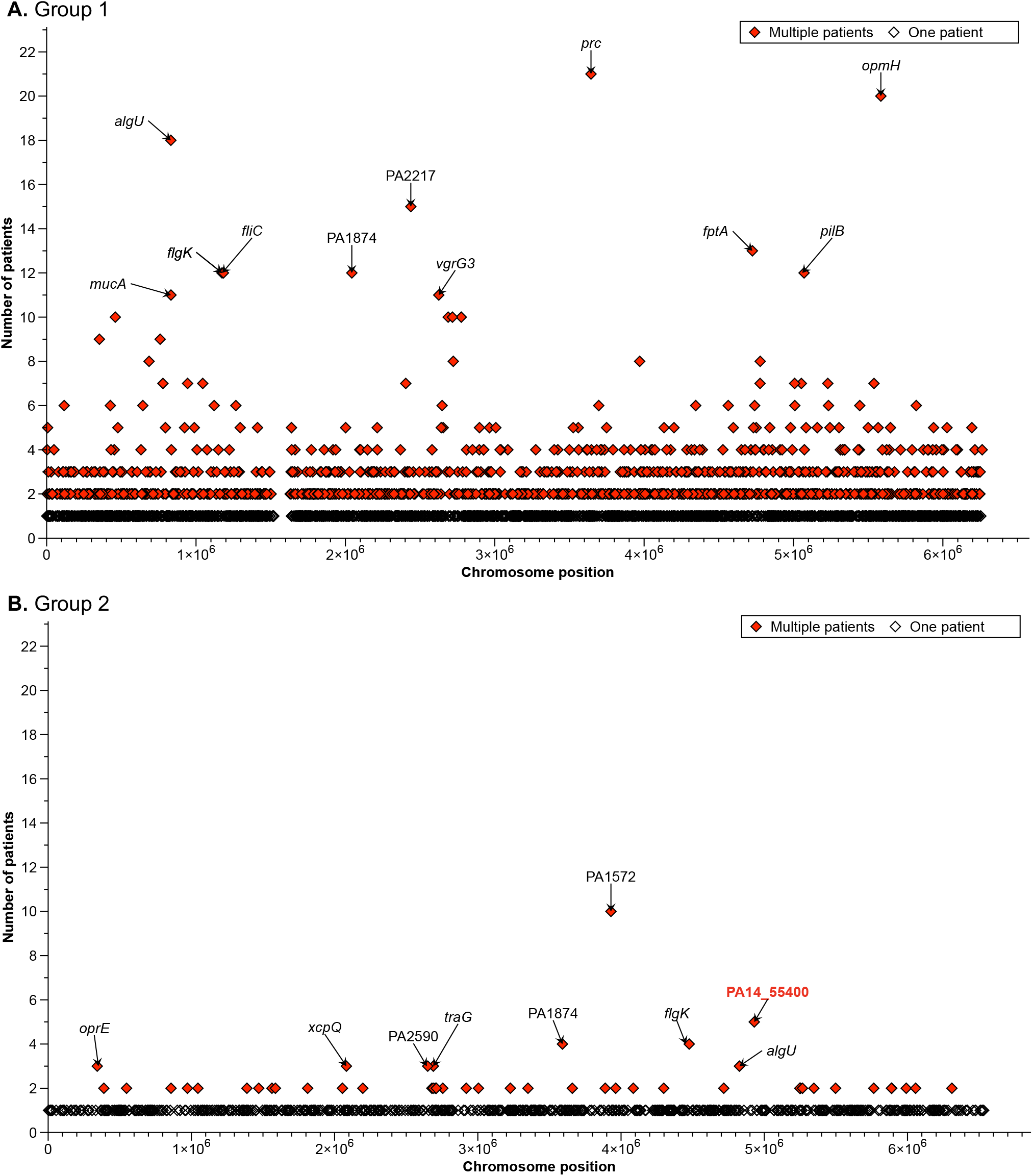
Within patient polymorphism. Each data point represents a gene with non-synonymous mutations between *Pseudomonas aeruginosa* isolates from the same bronchiectasis patient. They are filled if a mutation in the gene separates isolates in more than one patient (see legend). The y axis shows the number of patients with mutations between their sequenced isolates in each gene. The dataset has been split into phylogenetic group 1 **(A)** and group 2 **(B)** as different reference strains were used for each (group 1: PAO1, group 2: PA14). The top genes are labelled. The PA14 locus tags have been converted to PAO1 where possible for group 2, the red highlighted gene is not present in PAO1.

We next determined putative targets of positive selection, potentially involved in adaptation to the lung environment, by identifying phylogenetically independent SNPs that had become fixed in multiple patients (i.e., parallel evolution). Focusing on high impact mutations (e.g. interruption of start/stop codons) likely to cause loss of gene function (LoF), we observed putative parallel evolved mutations affecting 135 genes in group 1 isolates (Fig 4A) and 60 genes in group 2 isolates (Fig 4B). These included *prtN* (PA0610) which positively regulates pyocin expression^51^, the alginate biosynthesis regulator *algU*, and the cell-surface receptor *tonB*, linked to iron uptake, biofilm formation and quorum sensing^52^. Additionally, we observed evidence for parallel evolution in several hypothetical genes of unknown function, including PA4577, which shares homology with the transcriptional regulator *traR* (Table 1), and PA4788 and PA3036, which share homology with several dehydratases and oxidoreductases respectively. A large chromosomal region with no core SNPs in either group 1 or 2 (Fig 4) indicated large deletions occurring in a proportion of isolates; similar deletions have previously been associated with increased AMR in *P. aeruginosa* clinical isolates^53^.

**Table 1.** The most frequently occurring loss of function single nucleotide polymorphisms (SNPs) across *Pseudomonas aeruginosa* phylogenetic groups 1 and 2, and gene information. The phylogenetic group shows the group of isolates that carried the loss of function mutation in the gene, the number of patients is the bronchiectasis patients with isolates from that group that had the mutation, and the predicted number of independent mutation events indicates the number of independent mutation events found on the phylogeny for each mutation. There is one gene that appeared in the most frequent in both groups (*prtN*).

| Gene name | Gene product | Loss of function mutation | Phylogenetic group | Number of patients | Predicted number of independent mutation events |
| --- | --- | --- | --- | --- | --- |
| PA4638 | Hypothetical protein | Stop lost | 1 | 149 | 1 |
| PA2691 | Hypothetical protein | R87* | 1 | 119 | 1 |
| *hocS* | Short-chain O-acylcarnitine hydrolase | E322* | 1 | 102 | 1 |
| *pilQ* | Type 4 fimbrial biogenesis outer membrane protein PilQ precursor | S578* | 1 | 85 | 1 |
| PA4577 | Hypothetical protein | Q77* | 1 | 43 | 15 |
| PA4788 | Hypothetical protein | Q40* | 1 | 41 | 22 |
| PA3991 | Hypothetical protein | Q138* | 1 | 39 | 27 |
| PA1355 | Hypothetical protein | Start lost | 1 | 21 | 9 |
| PA3036 | Hypothetical protein | Y220* | 1 | 21 | 10 |
| *prtN* | Transcriptional regulator | Start lost | 1 | 20 (group 1) | 16 (group 1) |
|  |  | Start lost | 2 | 17 (group 2) | 1 (group 2) |
| *exoT* | Exoenzyme T | Stop lost | 2 | 25 | 1 |
| PA4062 | Hypothetical protein |  | 2 | 25 | 1 |
| *cat* | Chloramphenicol acetyltransferase | Stop lost | 2 | 25 | 1 |
| *waaC* | Heptosyltransferase I | Stop lost | 2 | 25 | 1 |
| PA14_00520 | Hypothetical protein | Stop lost | 2 | 21 | 1 |
| PA3577 | Hypothetical protein | Stop lost | 2 | 20 | 1 |
| *tonB* | TonB1 (part of TonB complex) | S306* | 2 | 18 | 1 |
| PA1606 | Hypothetical protein | S152* | 2 | 15 | 3 |
| PA5318 | Hypothetical protein | Stop lost | 2 | 15 | 1 |

**Figure 4.**
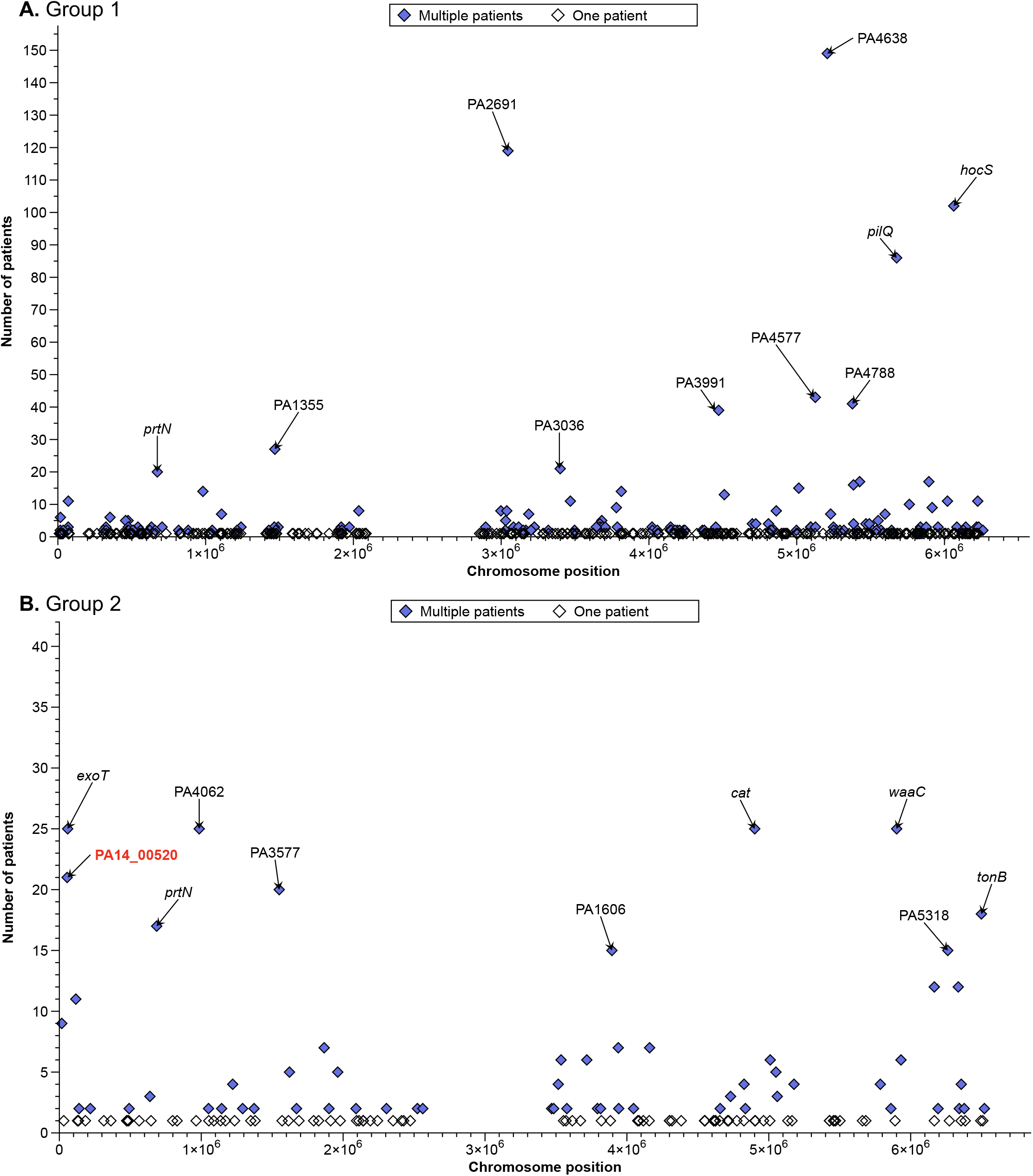
The number of bronchiectasis patients with at least one *Pseudomonas aeruginosa* isolate with a high impact single nucleotide polymorphism (SNP) (likely causing loss of function) in a gene. Each gene is represented by a data point. Group 1 **(A)** and group 2 **(B)** are shown on separate graphs as different reference strains were used for each (group 1: PAO1, group 2: PA14). The mutations in multiple patients are shown in blue, and the 10 most frequent amongst patients are labelled. The PA14 locus tags have been converted to PAO1 where possible, the red highlighted gene is not present in PAO1.

The abundance and distribution of AMR determinants within and between *P. aeruginosa* infections is poorly understood in bronchiectasis. Using the ResFinder database and the Comprehensive Antibiotic Resistance Database (CARD), we identified 16 variable AMR genes and 11 AMR-associated mutations across all isolates (Fig 5). Most were either fixed or absent in all isolates per patient, with rare occurrences of within patient polymorphism. Among the variable AMR genes likely to represent gain events, the most common was the ciprofloxacin modifying enzyme *crpP*, present in 49% of patients (Fig 5A). Other probable AMR gene gains were present in only one or a few patients, such as the aminoglycoside nucleotidyltransferase gene *aadA6* seen in 3 patients. These cases highlight the potential for the emergence of multidrug resistance in bronchiectasis. For example, 6 isolates from a single patient had an OXA-10 like beta-lactamase and an aminoglycoside acetyltransferase gene *aac(6’)-Ib-Hangzhou*, typically seen in *Acinetobacter baumanii* (Fig 5A)^54^, alongside the full complement of *P. aeruginosa* core AMR genes and *crpP*. AMR associated SNPs were found to commonly affect regulators of multidrug efflux systems, including *nalC, mexS* and *mexR*, and would be likely to cause upregulation of multidrug efflux (Fig 5B). Additionally, several common mutational targets were associated with resistance to fluoroquinolones, including L71R in the response regulator *pmrA* that was present in >80% of patients, and mutations in *gyrA* (T83I in 124 patients, and D87N in 14 patients) and *parE* (A473V in 14 patients), encoding topoisomerases targeted by ciprofloxacin. Whereas the *gyrA* mutations were mutually exclusive, the *parE* mutation co-occurred with the *gyrA* mutation T83I in 2 patients (Fig 5B).

**Figure 5.**
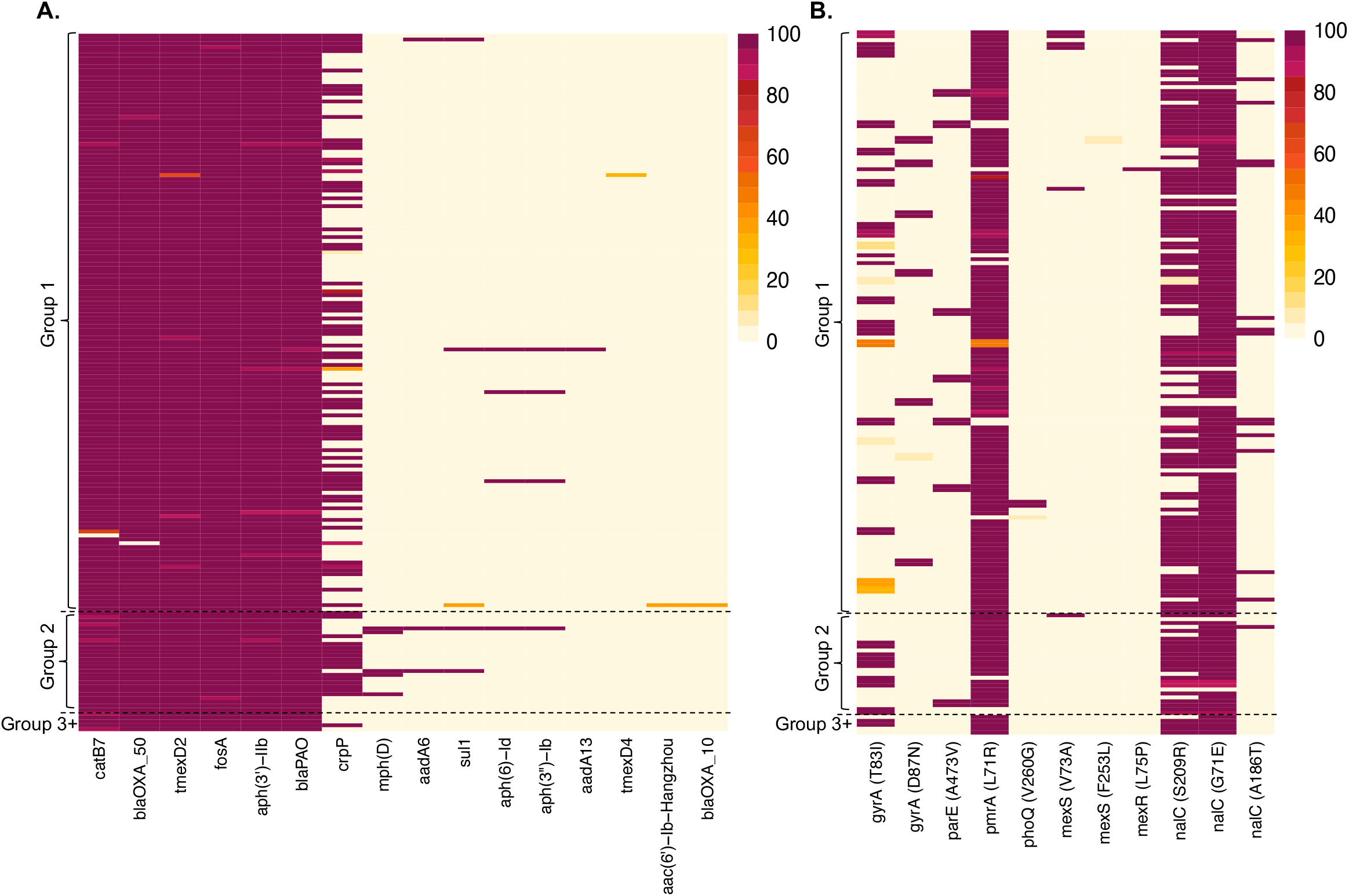
**(A)** The presence/absence of antimicrobial resistance (AMR) genes in each bronchiectasis *Pseudomonas aeruginosa* isolate identified using ResFinder. The patient without any OXA-50 beta-lactamases detected was found to have a large deletion in all isolates in this region (Fig S4). **(B)** The presence of AMR-associated mutations in each isolate based on Comprehensive Antibiotic Resistance Database (CARD) predictions, identified using RGI. In both heatmaps, each row represents a patient, and the fill colour shows the percentage of isolates from each patient that has the gene or mutation (see keys). Groups shown are the phylogenetic groups.

## Discussion

*P. aeruginosa* is the most common cause of respiratory infection in bronchiectasis worldwide, contributing to higher morbidity and mortality rates, but the genomic diversity of such infections is poorly understood. Our study provides a 10-fold increase in availability of genomic data for such infections, whilst also expanding patient sampling beyond Europe. Most infections were caused by a single, distinct ST and we found low rates of mixed ST infections and low incidence of CF epidemic clones, as well as little evidence of geographic variation in the causal STs. Accordingly, genetic diversity was higher between than within patients, but we nonetheless observed strong evidence for recurrent *in situ* diversification. In addition, we identified genes and functions undergoing parallel evolution independently in multiple patients associated with adaptation to the bronchiectasis lung environment. Although some of these functions, such as motility, biofilm and antimicrobial resistance, overlap with those commonly mutated in CF infections, others appear more bronchiectasis-specific, including several genes involved in pyocin production and a novel efflux pump.

Our findings provide strong evidence that *P. aeruginosa* undergoes evolutionary diversification and adaptation to the bronchiectasis lung environment in common with CF. Most patients are colonized with a single ST that diversifies *in situ* through both accumulation of mutations and changes in mobile genetic elements, predominantly changes in prophage number and gene content. Our evolutionary analysis revealed a suite of pathways recurrently under selection in multiple patients. Some functions gaining mutations are common to both bronchiectasis and CF, including regulators of alginate production (*mucA* and *algU*) associated with mucoidy and biofilm, flagellum and type-IV pilus genes (*flgK, fliC* and *pilQ*) likely to cause loss of motility, and genes involved in iron acquisition (including *tonB* and *fptA*) and cell envelope integrity (including *oprE, opmH* and *waaC*). In contrast, other mutational targets appear to be more common in bronchiectasis than in CF. These included mutations affecting pyocin production and resistance genes, perhaps suggesting that bacteriocin-mediated interference competition may be less intense compared to CF, and a novel efflux pump gene, PA1874, which was among the most frequently mutated in our patients but is not commonly mutated in CF infections. Mutations in this efflux pump have been linked to increased tobramycin resistance during biofilm infection^55^. Our finding of distinct evolutionary paths in bronchiectasis suggests that, despite some shared features with CF, there exist potentially important differences.

Initial colonization by *P. aeruginosa* in bronchiectasis is often treated with antibiotic regimens that include fluoroquinolone antibiotics^56^. Consistent with this, we commonly observed fluoroquinolone resistance genes (*crpP*) or mutations in targeted topoisomerases (*gyrA* and *parE*) or regulatory genes (*pmrA*) associated with resistance. After becoming chronically infected, bronchiectasis patients often receive long-term suppressive antibiotic therapy as well as antibiotic treatment for exacerbations, thus infections potentially receive treatment with a diversity of antibiotic classes. Accordingly, we commonly observed regulatory mutations likely to cause upregulation of multidrug efflux systems. This included mutation of a novel efflux pump (PA1874) as well as mutations affecting the MexAB-OprM that have not been highlighted previously in CF infections, potentially suggesting bronchiectasis-specific resistance mechanisms.

Consistent with prior single-country studies, our global genomic analysis revealed a high diversity of STs causing bronchiectasis infections across patients. Although we cannot distinguish the extent to which acquisition of infection is from patient-to-patient or environmental routes, our data clearly show that – unlike CF – highly-transmissible strains play only a minor role in the epidemiology of bronchiectasis infection. Unlike a previous UK study, we did not observe high incidence of mixed ST infections in our far larger sample of bronchiectasis infections which was each studied in greater depth than any previous study. Indeed, the very low rate of mixed ST infections alongside the low prevalence of CF epidemic STs implies that existing cohorting and isolation procedures for bronchiectasis are broadly effective at preventing dissemination of and superinfection by transmissible clones.

Owing to the serious negative impact of *P. aeruginosa* on patient health, improving treatment of such infections in bronchiectasis patients is a high priority. This study provides an unprecedented global genomic resource improving our knowledge and understanding of *P. aeruginosa* genetic diversity and evolution in bronchiectasis. Overall, our findings suggest important differences between bronchiectasis and CF infections, notably the relatively minor role that transmissible strains play in bronchiectasis relative to CF infections, and genes in the *P. aeruginosa* genome that are targets of selection in bronchiectasis but not CF.

## Acknowledgements

The ORBIT3 clinical trial was sponsored by Aradigm Corporation and samples kindly gifted to the University of Dundee and the European Bronchiectasis Network (EMBARC). We acknowledge the patients and investigators in the ORBIT programme. We thank all those involved in the clinical trial, including patients and all hospital and clinical trial staff, for the samples used for this work, as well as the Centre for Genomics Research (CGR) at the University of Liverpool for all sequencing.

## CRediT Statement

NEH: Methodology; Software; Formal analysis; Validation; Investigation; Data Curation; Writing -Original Draft; Writing - nReview & Editing; Visualization

AK: Methodology; Validation; Investigation; Data Curation; Writing - Review & Editing KC: Methodology; Validation; Investigation; Writing - Review & Editing

MJS: Investigation; Writing - Review & Editing

EMG: Methodology; Investigation

TF: Investigation RH: Data Curation

DZC: Conceptualization; Writing - Review & Editing; Funding acquisition

JLF: Conceptualization; Methodology; Resources; Writing - Original Draft; Writing - Review & Editing; Supervision

JDC: Conceptualization; Methodology; Resources; Writing - Original Draft; Writing - Review & Editing; Supervision; Funding acquisition

MAB: Conceptualization; Methodology; Resources; Writing - Original Draft; Writing - Review & Editing; Supervision; Project administration; Funding acquisition

SP: Conceptualization; Methodology; Software; Writing - Original Draft; Writing - Review & Editing; Resources; Supervision; Funding acquisition

