## Supplementary Information for "Global genomic diversity of *Pseudomonas aeruginosa* in bronchiectasis"

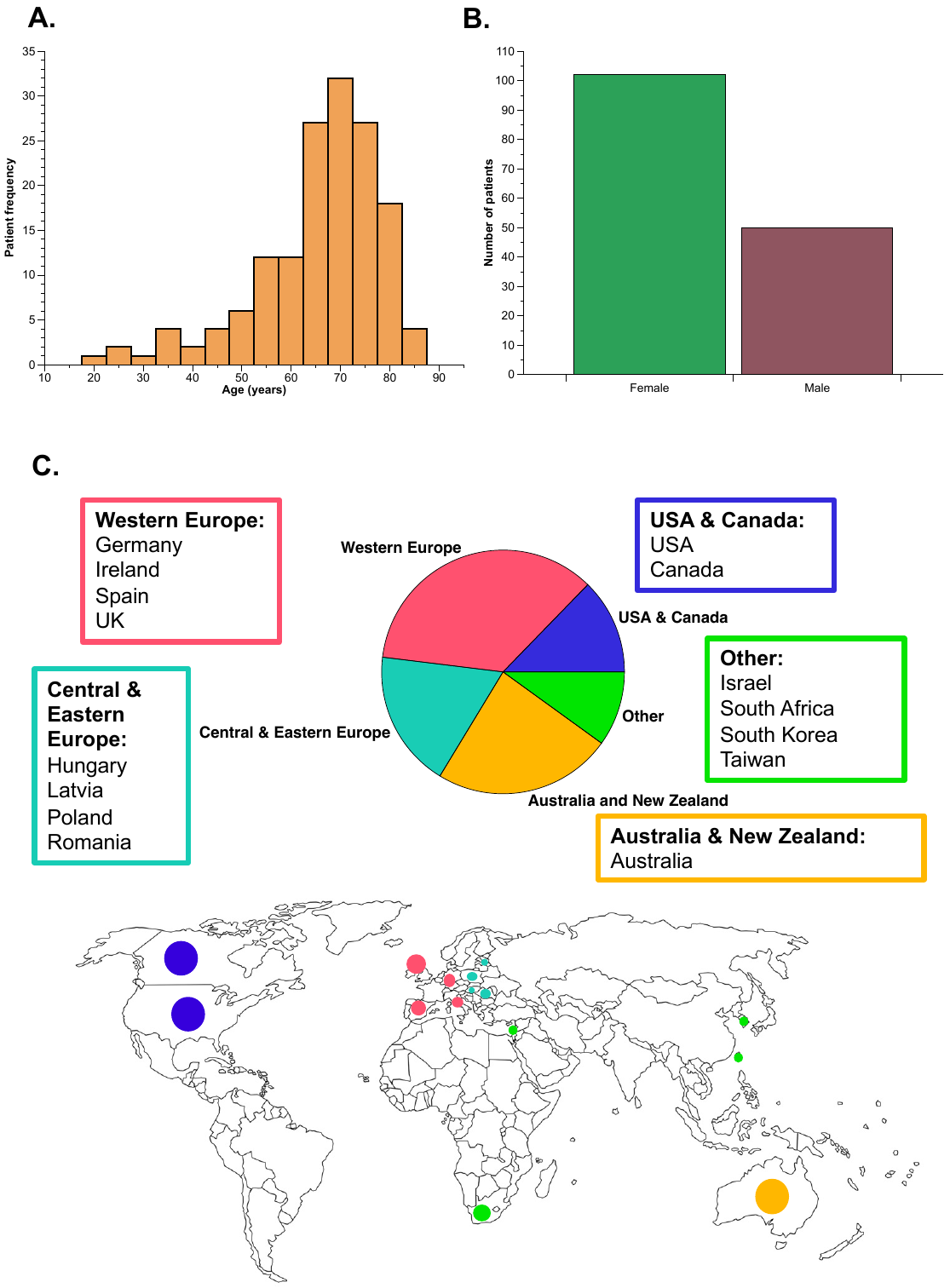

**Figure S1.** The demographic of bronchiectasis patients that sequenced *Pseudomonas aeruginosa* isolates were obtained from as part of the ORBIT3 clinical trial. **(A)** The distribution of ages at the start of the trial for all patients with available data. **(B)** The number of patients of each sex from available data. **(C)** The global regions represented by our sample set. The pie chart shows the proportion of patients attending clinics in each region, with the specific countries involved listed. The map shows these countries.

**Table S2.** The proportion of *Pseudomonas aeruginosa* isolates sequenced from people with bronchiectasis belonging to each phylogenetic group. Those in groups other than 1 and 2 have been grouped together based on their closest reference strain PA7, which is part of group 3, referred to as group 3+.

| **Phylogenetic group** | **Number of isolates** | **Patients (% (n))** |
| --- | --- | --- |
| 1 | 2359 | 82.78% (149) |
| 2 | 415 | 14.44% (26) |
| 3+ | 80 | 2.78% (5) |

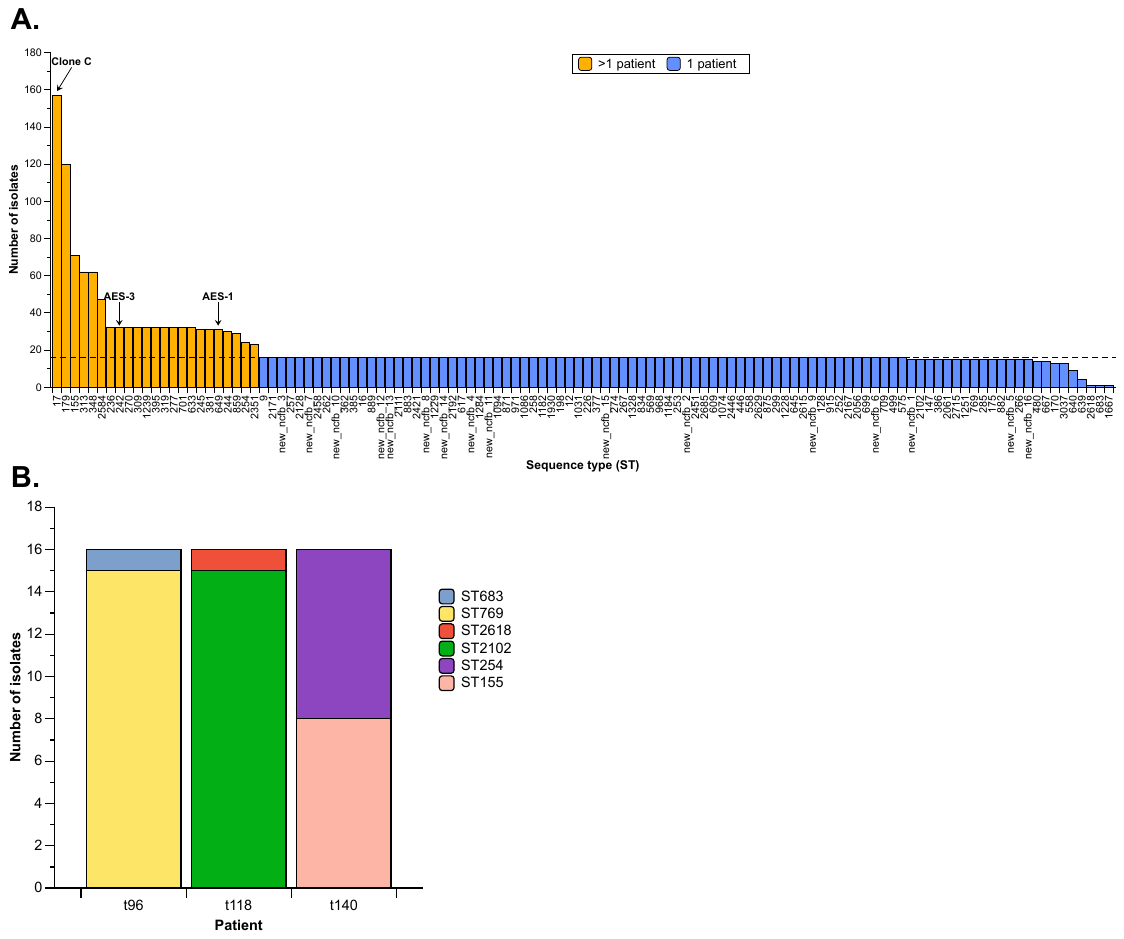

**Figure S2. (A)** *Pseudomonas aeruginosa* sequence types (STs) identified from multi-locus sequence type (MLST) analysis of isolates from people with bronchiectasis. The isolate frequency of all confirmed STs, both known and new (as labelled), is shown. The dashed line indicates 16 isolates, which was the total sequenced from each patient. Bars are colour coded based on whether the ST was present in multiple patients (see legend), those above the dashed line were present in >1 patient. Known STs and cystic fibrosis (CF) epidemic STs in multiple patients are labelled (AES: Australian Epidemic Strain). **(B)** Patients with more than one *P. aeruginosa* ST represented in the 16 isolates sequenced, indicating mixed-strain infection.

**Table S3.** The frequency of cystic fibrosis (CF) epidemic and prevalent sequence types (> 2 patients), including clone C, amongst sequenced *Pseudomonas aeruginosa* isolates from people with bronchiectasis, and the global region they were sampled from.

|  | **Sequence type (ST)** | **Number of isolates** | **Number of patients** | **Percentage of patients (n = 180)** | **Region** |
| --- | --- | --- | --- | --- | --- |
| **Frequently isolated** | | |  |  |  |
|  | 179 | 120 | 8 | 4.44% | Western Europe, Australia & New Zealand, USA & Canada, Other |
|  | 155 | 71 | 5 | 2.78% | Western Europe, Australia & New Zealand, Other |
|  | 313 | 62 | 4 | 2.22% | USA & Canada, Central & Eastern Europe, Other |
|  | 348 | 62 | 4 | 2.22% | Australia & New Zealand, USA & Canada |
|  | 2584 | 47 | 3 | 1.67% | Central & Eastern Europe |
| **Clonal/epidemic strains** | |  |  |  |  |
| **Clone C** | 17 | 157 | 10 | 5.56% | Western Europe, Australia & New Zealand, USA & Canada |
| **Liverpool epidemic strain (LES)** | 683 | 1 | 1 | 0.56% | Western Europe |
| **PA14-like** | 253 | 16 | 1 | 0.56% | Central & Eastern Europe |
| **Australian epidemic strain-1 (AES-1)** | 649 | 31 | 2 | 1.11% | Central & Eastern Europe, Other |
| **Australian epidemic strain-3 (AES-3)** | 242 | 32 | 2 | 1.11% | Australia & New Zealand |
| **DK-2** | 386 | 15 | 1 | 0.56% | Western Europe |
| **CC274** | 274 | 16 | 1 | 0.56% | Western Europe |

**Table S4.** The proportion of prophage regions detected amongst sequenced *Pseudomonas aeruginosa* isolates for each bronchiectasis patient using VirSorter. Each row represents a patient, with the patient code assigned in the first column. The remaining columns show the percentage of isolates sequenced with each number of prophage regions (see column headings). Empty boxes show where there are no isolates from the patient with that number of prophage regions. The percentages are colour coded from highest percentage (dark green) to lowest (light green).

|  | **Number of prophage regions** | | | | | | | | |
| --- | --- | --- | --- | --- | --- | --- | --- | --- | --- |
| **Patient** | **1** | **2** | **3** | **4** | **5** | **6** | **7** | **8** | **9** |
| t49 | **100** |  |  |  |  |  |  |  |  |
| t133 | **100** |  |  |  |  |  |  |  |  |
| p79 | **100** |  |  |  |  |  |  |  |  |
| p14 | **100** |  |  |  |  |  |  |  |  |
| t136 | **100** |  |  |  |  |  |  |  |  |
| p43 | **100** |  |  |  |  |  |  |  |  |
| t147 | **100** |  |  |  |  |  |  |  |  |
| p74 | **100** |  |  |  |  |  |  |  |  |
| t86 | **100** |  |  |  |  |  |  |  |  |
| t82 | **100** |  |  |  |  |  |  |  |  |
| p46 | **100** |  |  |  |  |  |  |  |  |
| t44 | **100** |  |  |  |  |  |  |  |  |
| t23 | **100** |  |  |  |  |  |  |  |  |
| p26 | **100** |  |  |  |  |  |  |  |  |
| t90 | **100** |  |  |  |  |  |  |  |  |
| t79 | **100** |  |  |  |  |  |  |  |  |
| t58 | **100** |  |  |  |  |  |  |  |  |
| t81 |  | **100** |  |  |  |  |  |  |  |
| t92 |  | **100** |  |  |  |  |  |  |  |
| p75 |  | **100** |  |  |  |  |  |  |  |
| t129 |  | **100** |  |  |  |  |  |  |  |
| t13 |  | **100** |  |  |  |  |  |  |  |
| t158 |  | **100** |  |  |  |  |  |  |  |
| t77 |  | **100** |  |  |  |  |  |  |  |
| t161 |  | **100** |  |  |  |  |  |  |  |
| p28 |  | **100** |  |  |  |  |  |  |  |
| t154 |  | **100** |  |  |  |  |  |  |  |
| p81 |  | **100** |  |  |  |  |  |  |  |
| t150 |  | **100** |  |  |  |  |  |  |  |
| t46 |  | **100** |  |  |  |  |  |  |  |
| p38 |  | **100** |  |  |  |  |  |  |  |
| p80 |  | **100** |  |  |  |  |  |  |  |
| t51 |  | **100** |  |  |  |  |  |  |  |
| p77 |  | **100** |  |  |  |  |  |  |  |
| p72 |  | **100** |  |  |  |  |  |  |  |
| t172 |  | **100** |  |  |  |  |  |  |  |
| p7 |  | **100** |  |  |  |  |  |  |  |
| t149 |  | **100** |  |  |  |  |  |  |  |
| t80 |  | **100** |  |  |  |  |  |  |  |
| t127 |  | **100** |  |  |  |  |  |  |  |
| p33 |  | **100** |  |  |  |  |  |  |  |
| p76 |  | **100** |  |  |  |  |  |  |  |
| t70 |  | **100** |  |  |  |  |  |  |  |
| t5 |  | **100** |  |  |  |  |  |  |  |
| p12 |  | **100** |  |  |  |  |  |  |  |
| t72 |  | **100** |  |  |  |  |  |  |  |
| t66 |  |  | **100** |  |  |  |  |  |  |
| t59 |  |  | **100** |  |  |  |  |  |  |
| p10 |  |  | **100** |  |  |  |  |  |  |
| t130 |  |  | **100** |  |  |  |  |  |  |
| t142 |  |  | **100** |  |  |  |  |  |  |
| t84 |  |  | **100** |  |  |  |  |  |  |
| t139 |  |  | **100** |  |  |  |  |  |  |
| p61 |  |  | **100** |  |  |  |  |  |  |
| p60 |  |  | **100** |  |  |  |  |  |  |
| t9 |  |  | **100** |  |  |  |  |  |  |
| t153 |  |  | **100** |  |  |  |  |  |  |
| t125 |  |  | **100** |  |  |  |  |  |  |
| t33 |  |  | **100** |  |  |  |  |  |  |
| p4 |  |  | **100** |  |  |  |  |  |  |
| t54 |  |  | **100** |  |  |  |  |  |  |
| t120 |  |  | **100** |  |  |  |  |  |  |
| t68 |  |  | **100** |  |  |  |  |  |  |
| t99 |  |  | **100** |  |  |  |  |  |  |
| p57 |  |  | **100** |  |  |  |  |  |  |
| t6 |  |  | **100** |  |  |  |  |  |  |
| t7 |  |  | **100** |  |  |  |  |  |  |
| t10 |  |  | **100** |  |  |  |  |  |  |
| t89 |  |  | **100** |  |  |  |  |  |  |
| t155 |  |  |  | **100** |  |  |  |  |  |
| t71 |  |  |  | **100** |  |  |  |  |  |
| t141 |  |  |  | **100** |  |  |  |  |  |
| t168 |  |  |  | **100** |  |  |  |  |  |
| p69 |  |  |  | **100** |  |  |  |  |  |
| p42 |  |  |  | **100** |  |  |  |  |  |
| p23 |  |  |  | **100** |  |  |  |  |  |
| t152 |  |  |  | **100** |  |  |  |  |  |
| p82 |  |  |  | **100** |  |  |  |  |  |
| t109 |  |  |  | **100** |  |  |  |  |  |
| t78 |  |  |  | **100** |  |  |  |  |  |
| t94 |  |  |  |  | **100** |  |  |  |  |
| t113 |  |  |  |  | **100** |  |  |  |  |
| t156 |  |  |  |  | **100** |  |  |  |  |
| t122 |  |  |  |  |  | **100** |  |  |  |
| p59 |  |  |  |  |  | **100** |  |  |  |
| t61 |  |  |  |  |  | **100** |  |  |  |
| p50 |  |  |  |  |  | **100** |  |  |  |
| t85 |  |  |  |  |  |  | **100** |  |  |
| t87 |  |  | **6** | **94** |  |  |  |  |  |
| p21 |  |  | **6** | **94** |  |  |  |  |  |
| p78 |  |  | **6** | **94** |  |  |  |  |  |
| t110 |  |  | **6** |  | **94** |  |  |  |  |
| p70 |  |  |  | **6** | **94** |  |  |  |  |
| p40 |  | **6** | **94** |  |  |  |  |  |  |
| t143 |  | **6** | **94** |  |  |  |  |  |  |
| t165 |  |  | **6** |  | **94** |  |  |  |  |
| t107 | **94** |  | **6** |  |  |  |  |  |  |
| p67 |  |  | **94** | **6** |  |  |  |  |  |
| t1 |  | **6** |  | **94** |  |  |  |  |  |
| t128 |  |  | **6** | **94** |  |  |  |  |  |
| t132 |  |  |  | **6** | **94** |  |  |  |  |
| t157 | **6** | **94** |  |  |  |  |  |  |  |
| t151 |  |  | **6** | **94** |  |  |  |  |  |
| p63 |  | **94** | **6** |  |  |  |  |  |  |
| t55 |  | **6** | **94** |  |  |  |  |  |  |
| t12 |  | **6** | **94** |  |  |  |  |  |  |
| p49 |  | **6** | **94** |  |  |  |  |  |  |
| t174 |  |  |  |  |  | **6** | **94** |  |  |
| t144 |  | **6** | **94** |  |  |  |  |  |  |
| t116 |  |  |  | **6** | **94** |  |  |  |  |
| t123 |  | **7** | **93** |  |  |  |  |  |  |
| t101 |  |  |  | **8** | **92** |  |  |  |  |
| t88 | **6** | **31** | **63** |  |  |  |  |  |  |
| p19 |  |  | **75** | **19** | **6** |  |  |  |  |
| p71 |  | **31** | **44** | **6** | **13** | **6** |  |  |  |
| p37 |  | **13** | **81** | **6** |  |  |  |  |  |
| t75 |  | **19** | **81** |  |  |  |  |  |  |
| t73 |  |  | **6** | **56** | **32** | **6** |  |  |  |
| t166 |  |  | **19** | **81** |  |  |  |  |  |
| t24 |  |  | **62** | **38** |  |  |  |  |  |
| p18 |  | **19** | **81** |  |  |  |  |  |  |
| t148 |  | **25** | **75** |  |  |  |  |  |  |
| t30 |  | **38** | **62** |  |  |  |  |  |  |
| p85 |  |  |  |  |  | **50** | **25** | **6** | **19** |
| t146 |  |  | **6** | **6** | **88** |  |  |  |  |
| t69 |  |  |  | **6** | **81** | **13** |  |  |  |
| p68 |  | **88** | **12** |  |  |  |  |  |  |
| t93 |  | **62** | **38** |  |  |  |  |  |  |
| t91 | **6** | **69** | **25** |  |  |  |  |  |  |
| p24 |  |  | **71** | **29** |  |  |  |  |  |
| t52 |  |  | **6** | **47** | **47** |  |  |  |  |
| p64 | **6** | **25** | **69** |  |  |  |  |  |  |
| p62 |  |  | **69** | **31** |  |  |  |  |  |
| p45 |  |  |  | **6** | **38** | **56** |  |  |  |
| t32 |  |  |  |  | **88** | **12** |  |  |  |
| t106 |  |  |  |  | **13** | **74** | **13** |  |  |
| t124 |  | **69** | **31** |  |  |  |  |  |  |
| p25 |  |  |  |  |  |  | **12** | **88** |  |
| p15 |  |  |  | **6** | **6** | **50** | **6** | **26** | **6** |
| t39 |  |  | **12** | **88** |  |  |  |  |  |
| p56 |  |  |  |  | **6** | **81** | **13** |  |  |
| t162 | **12** | **88** |  |  |  |  |  |  |  |
| t145 |  |  |  | **81** | **6** | **13** |  |  |  |
| p9 |  |  | **56** | **44** |  |  |  |  |  |
| t159 |  | **6** | **56** | **38** |  |  |  |  |  |
| t95 |  |  |  |  | **6** |  | **69** | **25** |  |
| t98 |  |  |  | **38** | **62** |  |  |  |  |
| T167 |  |  |  | **31** | **69** |  |  |  |  |
| p84 | **38** | **62** |  |  |  |  |  |  |  |
| t36 |  | **62** | **38** |  |  |  |  |  |  |
| p16 |  | **19** | **81** |  |  |  |  |  |  |
| p39 |  | **12** | **88** |  |  |  |  |  |  |
| p6 |  |  |  | **56** | **44** |  |  |  |  |
| t160 |  | **19** | **81** |  |  |  |  |  |  |
| p17 |  | **6** | **88** | **6** |  |  |  |  |  |
| t112 |  |  |  | **13** | **7** | **80** |  |  |  |
| t111 |  | **69** | **31** |  |  |  |  |  |  |
| t117 | **81** | **19** |  |  |  |  |  |  |  |
| t169 | **88** | **12** |  |  |  |  |  |  |  |
| t48 |  | **50** | **44** | **6** |  |  |  |  |  |
| p36 |  | **31** | **69** |  |  |  |  |  |  |
| p48 |  |  |  | **75** | **25** |  |  |  |  |
| t108 |  |  |  | **88** | **12** |  |  |  |  |
| t3 |  |  | **12** | **88** |  |  |  |  |  |
| t63 |  |  | **31** | **50** | **19** |  |  |  |  |
| t16 |  |  | **56** | **38** | **6** |  |  |  |  |
| t114 |  | **56** | **44** |  |  |  |  |  |  |
| t83 |  |  | **26** | **31** | **31** | **6** | **6** |  |  |
| t100 |  |  |  | **75** | **25** |  |  |  |  |
| p20 |  |  |  | **13** | **31** |  | **56** |  |  |
| t74 |  |  |  | **12** | **88** |  |  |  |  |
| p55 |  |  |  | **12** | **44** | **44** |  |  |  |
| t62 |  |  |  | **12** | **88** |  |  |  |  |
| p22 |  |  | **12** | **88** |  |  |  |  |  |
| p58 |  | **6** | **25** | **69** |  |  |  |  |  |
| t97 |  |  | **6** | **6** | **88** |  |  |  |  |
| p51 |  |  |  | **6** | **88** | **6** |  |  |  |

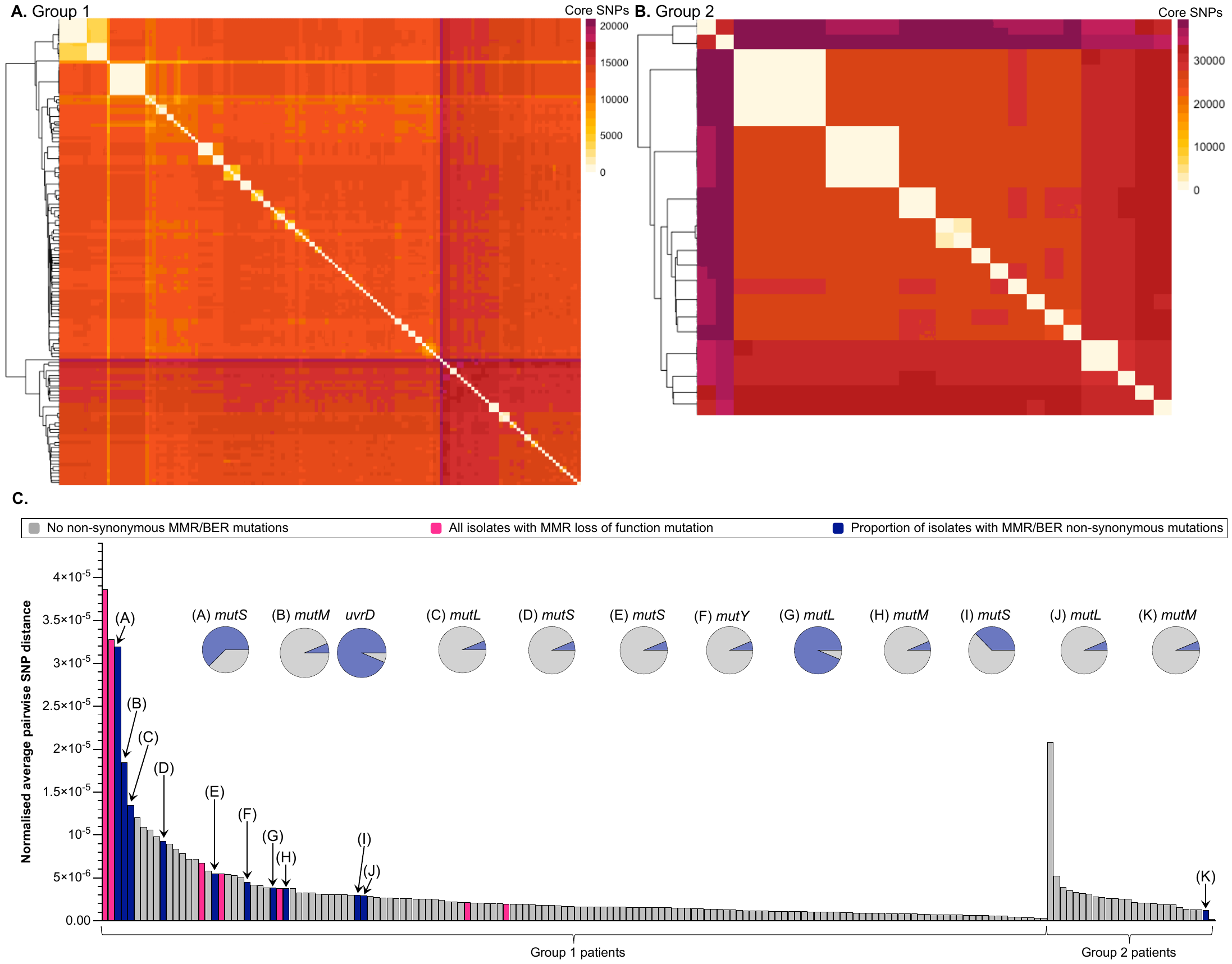

**Figure S3. (A,B)** Heatmaps showing the core single nucleotide polymorphism (SNP) pairwise distance matrix of phylogenetic group 1 (A) and phylogenetic group 2 (B) *Pseudomonas aeruginosa* bronchiectasis isolates. The fill represents the number of core SNPs between each isolate (see legend), and the light coloured diagonal line shows same patient isolate comparisons. **(C)** The normalised average pairwise SNP distance for each patient, calculated as the average pairwise core SNP distance between isolates from the same patient divided by the number of core nucleotides. Groups shown are phylogenetic groups. The patients with all isolates carrying a loss of function mutation in DNA mismatch repair (MMR) genes are highlighted in pink. No loss of function mutations in base excision repair (BER) genes were detected. Patients with non-synonymous mutations between isolates (i.e. polymorphic) are labelled with letters corresponding to pie charts that show the proportion of sequenced isolates carrying the mutation (blue = with mutation, grey = without).

**Table S5.** The genes that most frequently carried non-synonymous single nucleotide polymorphisms (SNPs) between *Pseudomonas aeruginosa* isolates from the same bronchiectasis patient amongst either phylogenetic group 1 (PAO1-like) or group 2 (PA14-like). The total number of patients shows number of patients across both groups that have polymorphism in the gene. The proportion of isolates column bar charts show the proportion of sequenced isolates per patient with a non-synonymous mutation in the gene; the blue shows isolates with a mutation (top bar) and the grey shows isolates without a mutation (bottom bar), the y axis is the number of isolates.

| **Gene name** | **Gene product** | **Total number of patients** | **Proportion of isolates** |
| --- | --- | --- | --- |
| *prc* | Periplasmic tail-specific protease | 21 | 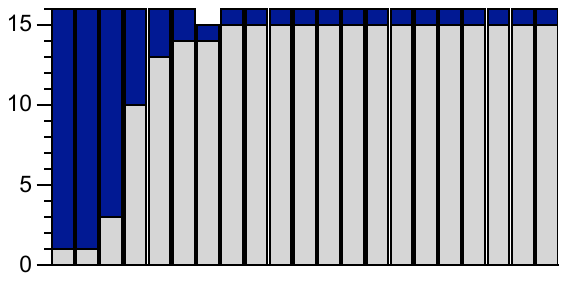 |
| *opmH* | Probable outer membrane protein precursor | 20 | 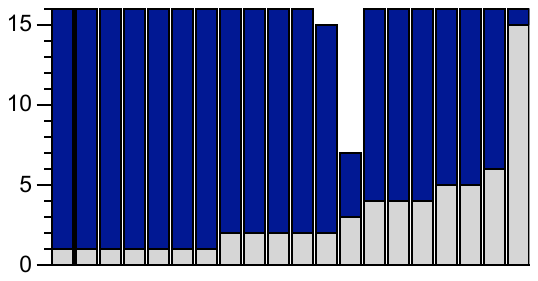 |
| *algU* | Sigma factor | 20 | 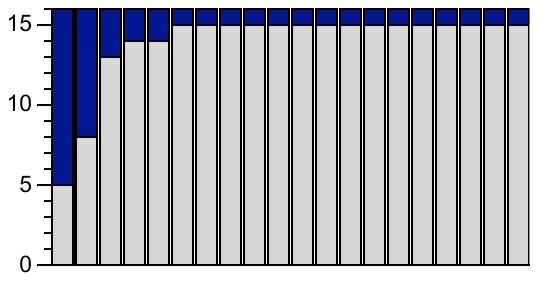 |
| PA2217 | Probable aldehyde dehydrogenase | 16 | 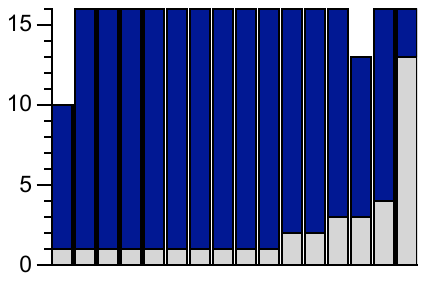 |
| *flgK* | Flagellar hook-associated protein 1 | 16 | 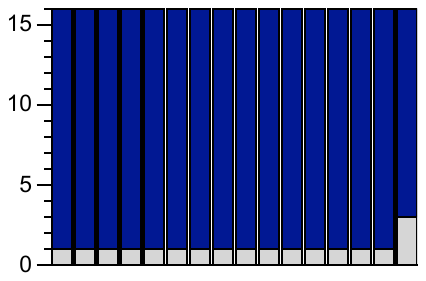 |
| PA1874 | Hypothetical protein | 16 | 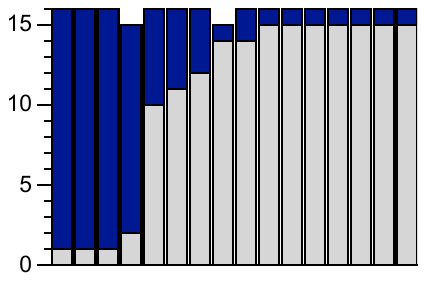 |
| *fptA* | Fe(III)-pyochelin outer membrane receptor precursor | 14 | 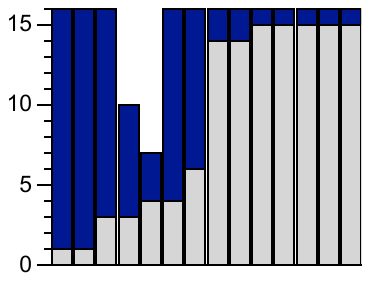 |
| *fliC* | Flagellin type B | 13 | 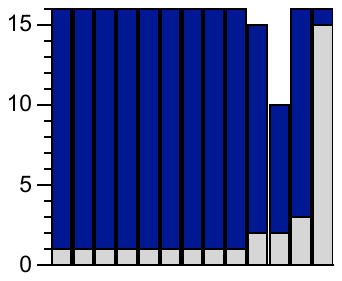 |
| *pilB* | Type 4 fimbrial biogenesis protein | 12 | 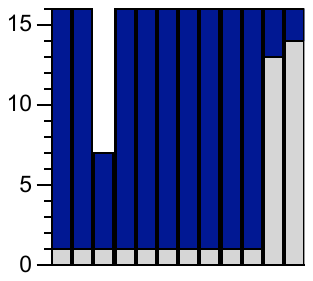 |
| *vgrG3* | VgrG3 | 12 | 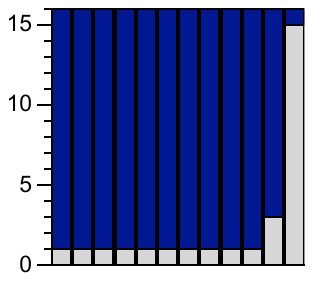 |
| *mucA* | Anti-sigma factor | 11 | 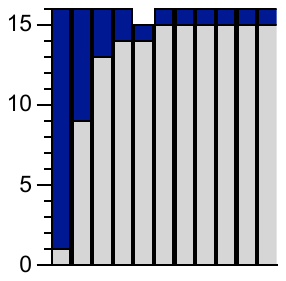 |
| PA1572 | Hypothetical protein | 11 | 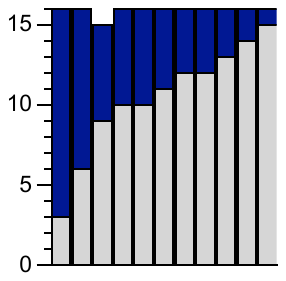 |
| *xcpQ* | General secretion pathway protein D | 6 | 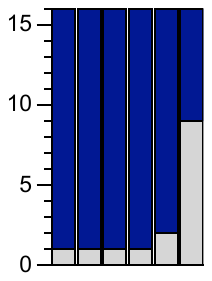 |
| PA14_55400 | Hypothetical protein | 5 | 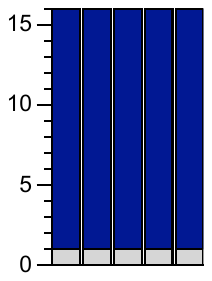 |
| *oprE* | Anaerobically-induced outer membrane porin OprE precursor | 5 | 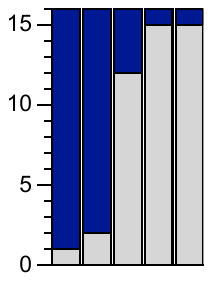 |
| PA2590 | Hypothetical protein | 5 | 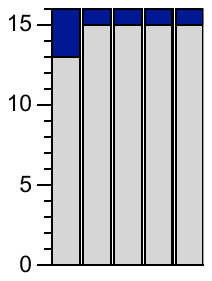 |
| *traG* | Conjugal transfer coupling protein | 3 | 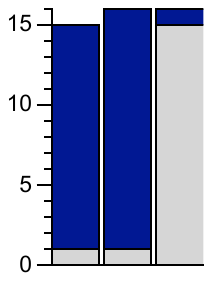 |

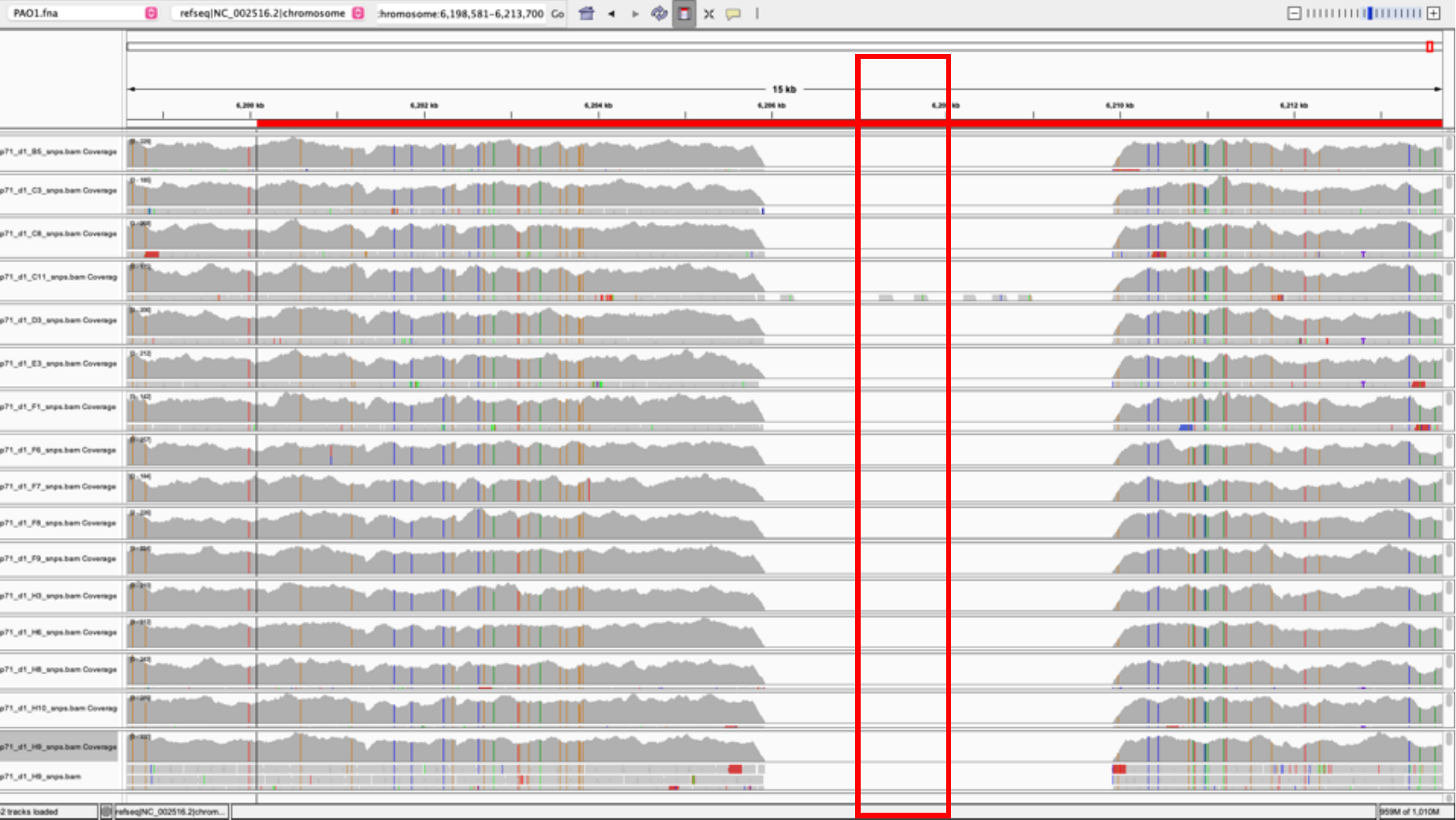
**Figure S4.** *Pseudomonas aeruginosa* isolates from the one bronchiectasis patient without an OXA-50-like beta-lactamase (region shown by the red box) due to the large deletion shown.
